# Adaptive Group Behavior of Fragile X Mice in Unfamiliar Environments

**DOI:** 10.1101/2023.09.26.559495

**Authors:** Gabriele Giua, Benjamin Strauss, Olivier Lassalle, Pascale Chavis, Olivier J. Manzoni

**Author notes:** Corresponding author: Olivier Manzoni, PhD., INMED, INSERM U901, Parc Scientifique de Luminy, BP13 – 13273 Marseille Cedex 09, France. These authors have contributed equally to this work (Gabriele Giua, Benjamin Strauss). Lead contact: Olivier Manzoni, PhD.

## Abstract

Fragile X Syndrome (FXS) stands out as a prominent cause of inherited intellectual disability and a prevalent disorder closely linked to autism. FXS is characterized by substantial alterations in social behavior, encompassing social withdrawal, avoidance of eye contact, heightened social anxiety, increased arousal levels, language deficits, and challenges in regulating emotions. Conventional behavioral assessments primarily focus on short-term interactions within controlled settings. In this study, we conducted a comprehensive examination of the adaptive group behavior of FXS mice over a three-day period, without introducing experimental interventions or task-based evaluations. The data unveiled intricate behavioral anomalies, with the most significant changes manifesting during the initial adaptation to unfamiliar environments. Notably, certain behaviors exhibited a gradual return to typical patterns over time. This dynamic FXS phenotype exhibited heightened activity, featuring increased exploration, amplified social interest, and an unconventional approach to social interactions characterized by a higher frequency of shorter engagements. These findings contribute to the growing understanding of social behavior in individuals with FXS and underscore the significance of comprehending their adaptive responses in various environmental contexts.

## Introduction

Fragile X Syndrome (FXS) stands as the primary cause of inherited intellectual disability (ID) and ranks as the most prevalent syndrome related to autism spectrum disorders (ASD)^1,2^. It affects approximately 1.4 in 10,000 males and 0.9 in 10,000 females^1^. FXS is the result of a mutation in the *FMR1* gene, which impedes the production of Fragile X Messenger Ribonucleoprotein 1 (FMRP), a crucial protein for neurodevelopment and the maintenance of neuronal and synaptic functions^3^. Insufficiency of FMRP leads to significant abnormalities in the central nervous system, contributing to the cognitive and behavioral disorders observed in individuals with FXS^4^. The FXS phenotype is characterized by significant alterations in social behavior, which profoundly impact the well-being of individuals and their families. These symptoms encompass social withdrawal^5–7^, avoidance of gaze^8–10^, social anxiety^11–13^, hyperarousal^14^, deficits in language development^15,16^, and difficulties in recognizing and regulating emotions^17,18^. These symptoms collectively contribute to the deterioration of the social competence of individuals with FXS.

Assessing mouse social behavior under both normal and pathological conditions is crucial for gaining insights into the neural systems implicated in psychiatric disorders. Many studies have used standardized short-term social interaction assessments, such as the 3-chamber test, to investigate the social interaction behavior of FXS mice^19–24^. The studies have reported various alterations in the social behavior of these mice, but the results have shown heterogeneity, likely attributable to differences in the protocols used^25^. Standardized tests primarily assess parameters related to simple, short-lived social interactions occurring in controlled, unfamiliar settings. Additionally, within the context of ASD, the experimenter manipulation and the unfamiliar settings typically encountered in task-based assessments can act as significant stressors, profoundly impacting animal behavior and hindering the observation of complex, naturalistic behaviors^26–28^. Furthermore, this approach lacks important ethological behavioral markers, such as group dynamics and long-term interactions^29^.

In this study, we leveraged the Live Mouse Tracker^30^, a tool that facilitates continuous, unsupervised, and non-invasive monitoring of spontaneous social interactions among groups of mice. Our objective was to explore the social behavioral traits associated with FXS within an environment designed to mimic their standard housing and enrichment conditions. This innovative approach removes the necessity for task-based assessments and allows for the analysis of group behaviors over extended time periods.

The data presented here unveils a spectrum of intricate anomalies within the exploratory and social behavior of FXS mice. The most pronounced variations were observed during their initial adaptation to the unfamiliar environment. However, some altered behaviors exhibited a gradual attenuation, eventually normalizing within a two-day period. This dynamic FXS phenotype manifests as a hyperactive profile, marked by a heightened propensity for exploring new surroundings, an increased interest in social interaction combined to a distinctive pattern of social engagements characterized by more frequent but shorter interactions in comparison to the control group.

### Methods

### Animals

Animals were treated in compliance with the European Communities Council Directive (86/609/EEC) and the United States National Institutes of Health Guide for the care and use of laboratory animals. The French Ethical committee authorized this project (APAFIS#34573-202201071122121v3). *Fmr1*-KO2 mice from FRAXA foundation were used in this study. Females *Fmr1+/-* were paired with males *Fmr1+/y, both with* C57Bl/6J background. Pups were weaned and ear punched for identification and genotyping at postnatal day 21 (P21). *Fmr1+/y* mice composed the control group (wild type (WT)) and *Fmr1-/y* the experimental group (knockout, KO). Between P30 and P35 mice were organized in experimental groups into cages of 4 males of the same genotype. Mice were housed in standard wire-topped Plexiglas cages (42□×□27□x□14□cm) in a temperature and humidity-controlled condition (i.e., temperature 21□±□1□°C, 60L±□10% relative humidity and 12h light/dark cycles). Food and water were available ad *libitum*.

### Live Mouse Tracker

To perform real-time behavior acquisition/analysis of group-housed mice the Live Mouse Tracker (LMT), leverages infrared tracking, RFID-chip identification, and machine learning^30^. The LMT arena is a 50×50×30cm cage made in transparent PMMA. For each LMT run, the arena was covered with 1kg of litter. Two squared houses (10×10×5cm) of red transparent PMMA were placed in two opposite corners of the arena. Two cotton rolls were placed next to each house. A U-shaped house (7.62×9.5×4.5cm) of red transparent PMMA was placed between the two squared houses for enrichment. 100g of standard chow (an ad *libitum* quantity for the experiment duration) was placed in the arena. Mice had *ad libitum* access to two bottles of drinking water placed in the two corners opposite to squared houses. The disposition of the elements in the arena is shown in Fig. 1A.

**Figure 1.**
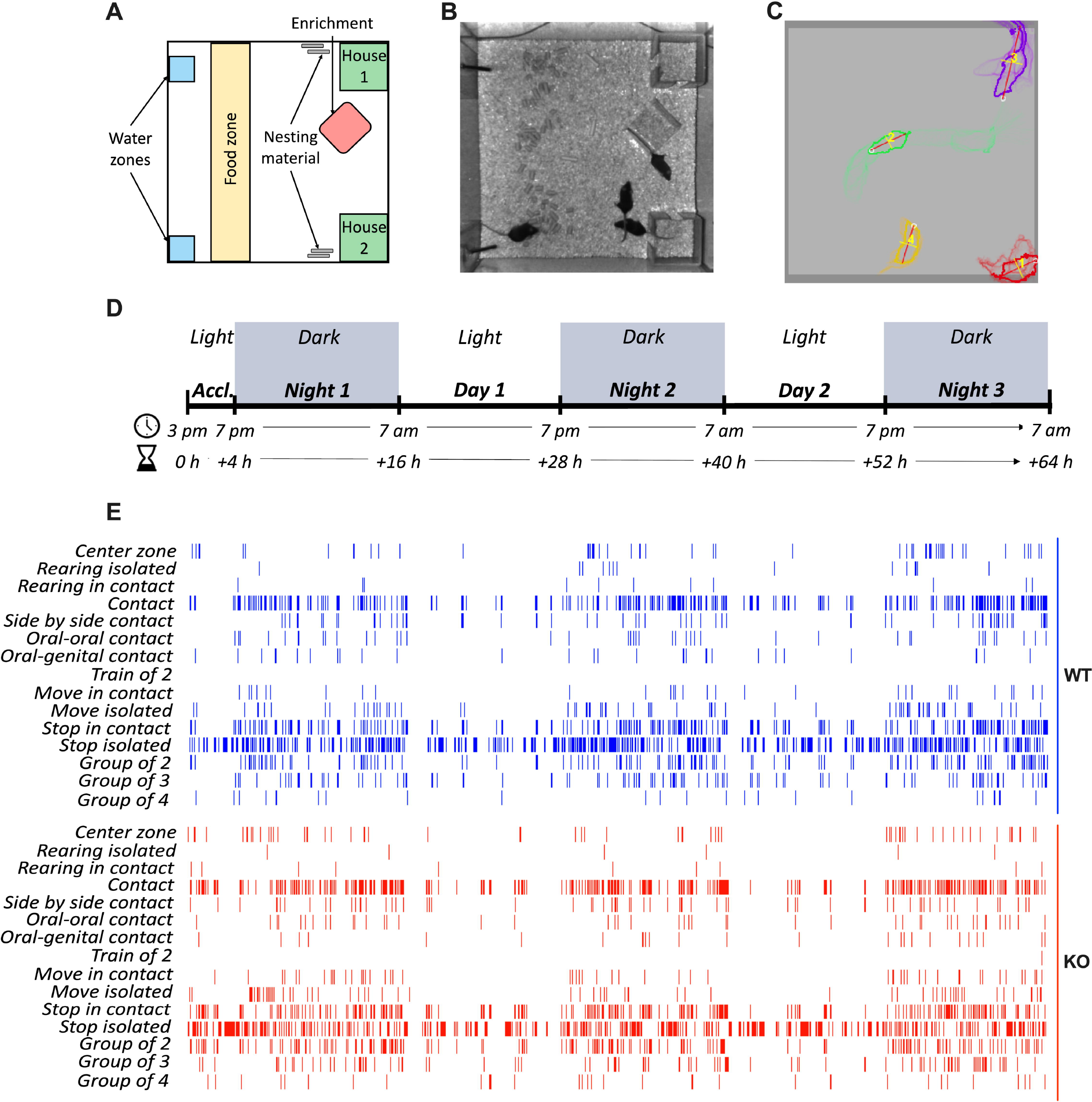
Experimental setting. **A**. Scheme of the arena showing 2 accesses to 2 water bottles (blue), 1 area representing the food distribution area (yellow), 2 houses (green), environmental enrichment (red) and nesting material (gray). **B**. Picture of the arena and a group of 4 mice during the experiment. **C**. Live tracking of the mice, representing the mask and trajectory of each. **D**. Experimental timeline representing the light/dark cycles over the duration of the experiment. **E**. Example of the detection of different behaviors of a single mouse throughout the entire experiment.

LMT behavioral experiments were performed on groups of 4 four male mice of the same genotype, aged between P70 and P90. Each group of mice was subjected to a single 64h LMT trial. Each trial began at 3 pm and included an initial 4h light period followed by three dark and two light alternated 12h-cycles (Fig. 1D). Sound, humidity, temperature, and luminosity were monitored during experiments. No access to the LMT room was allowed during trial. Mice weight, food and water consumption were monitored before and after the experiment.

### RFID

Adolescent male mice (P30-35) were deeply anesthetized with isoflurane and subcutaneously injected with a RFID chip (Biomark, APT12) using Biomark MK25 PIT Tag, behind the ears of the animal, and gently pushed to the back of the animal. The RFID implantation was performed minimum 35 days before the LMT trial to avoid injection-induced stress impact on mouse behavior.

### Data analysis

During each trial, the movements of each mouse within a social group of 4 individuals were tracked capturing 30 frames per second. LMT data were extracted from the SQLite database using scripts from either the LMTAnalysisMaster packages or newly created scripts available on GitLab (https://gitlab.com/chavis_manzoni_lab/lmt-scripts), both utilized with Pydev in Eclipse. Several types of behaviors were extracted from the dataset (list and descriptions is provided in supplementary information in De Chaumont et al., 2019^30^).

### Statistics and graphs

With the exception of Principal Component Analysis (PCA, see below), statistical analysis and graphic representation of data were performed with Prism (GraphPad Software). Statistical comparison was based on nonparametric Mann-Whitney U test. Considering the numerous parameters reported in the evaluation of behavior, PCA was computed using FactoMineR package^31^ with R (RCore Team (2021). R: A language and environment for statistical computing. R Foundation for Statistical Computing, Vienna, Austria. URL https://www.R-project.org/).

The animal illustrations featured in the graphs were created using BioRender (URL https://www.biorender.com)

### Code availability

Newly created scripts available on GitLab (https://gitlab.com/chavis_manzoni_lab/lmt-scripts).

## Results

The activity of *Fmr1* WT or KO mice was studied within homogeneous groups of four mice of the same sex and genotype in an arena that mimicked their housing conditions (Fig. 1A–C). The recordings covered a continuous 64-hour period, starting with a 4-hour adjustment phase to the new environment. Afterward, there was a cycle of three 12-hour nocturnal phases followed by two 12-hour diurnal phases, as shown in Fig. 1D. Throughout these phases, every single mouse within the arena was continuously monitored along the x, y, and z axes, enabling the construction of a comprehensive behavioral profile based on the detection of various exploratory and social parameters (Fig. 1E).

### Normal day/night rhythm in groups of FXS mice

Activity in our group of mice was monitored across alternating light and dark phases (Fig. 2). Similar temporal dynamics of locomotor activity were observed in both genotypes (Fig. 2B). There was a significant surge in locomotor activity within the initial hour during the initial phase of the experiment when mice were first introduced to the new environment for acclimation. This heightened activity gradually diminished over the following three hours leading up to the onset of the first dark phase. Throughout all observed dark cycles, a uniform activity rhythm became evident. Over these 12-hour periods, there was a noticeable spike in activity during the initial 4 hours, followed by a 4-hour period of reduced activity, and finally, a secondary activity peak (although smaller than the initial one) in the last 4 hours of the night. As expected, daytime cycles were characterized by low levels of mouse activity, occasionally punctuated by minor activity spikes (Fig. 2B). Based on the time course of motor activity, we selected three specific time points for further in-depth quantitative and qualitative analysis of exploratory and social behaviors. These time points corresponded to the highest peaks of activity, representing periods during which exploratory behavior and active social interaction among cagemates were most pronounced.

**Figure 2.**
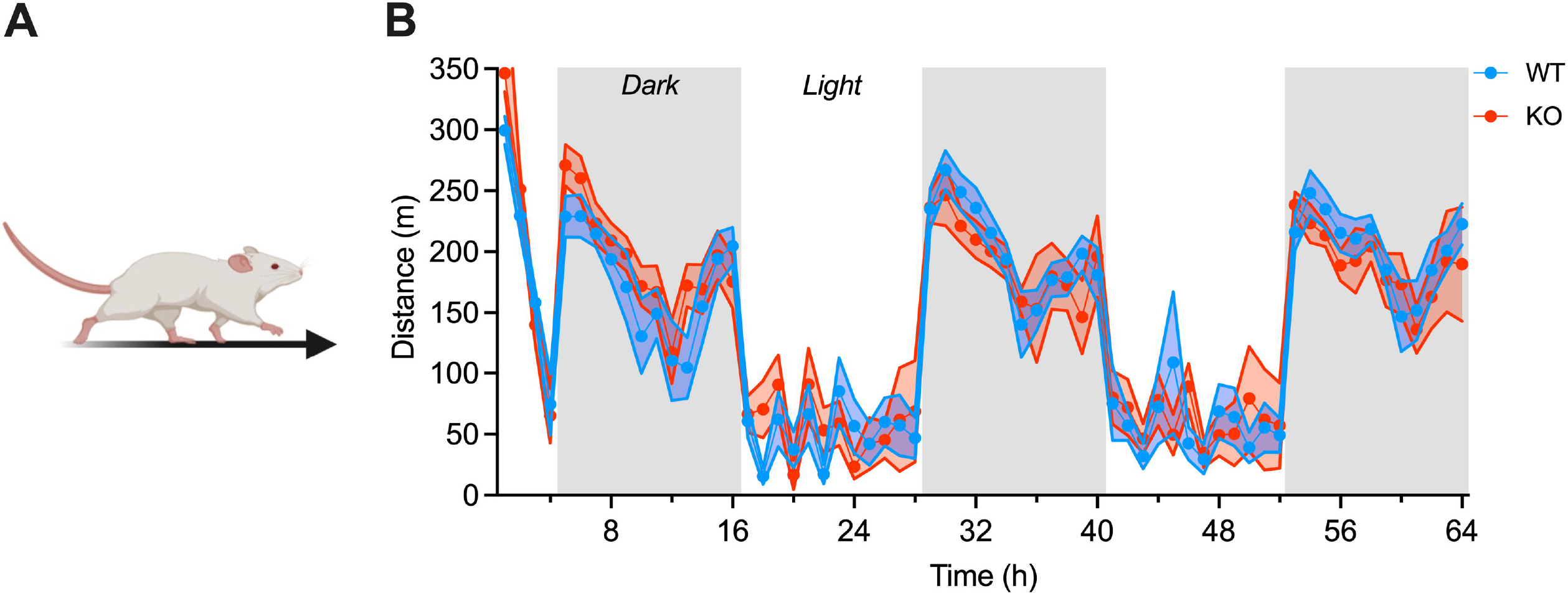
Locomotor activity day/night temporal dynamics are normal in FXS mice. **A**. Illustration of a mouse in motion. **B**. The temporal dynamics of activity, measured as distance traveled, traced a similar day/night rhythm between Fmr1 WT and KO mice. Single dot represents group mean value at that current step. Data are shown as mean ± 95% CI in XY plot. WT group in blue, N=20; KO group in orange, N=20.

Specifically, these time points coincided with 1/ the peak of activity during the first hour in the arena (i.e., 0-1h, Fig. 2B); 2/ the peak of activity during the first 4 hours of the first dark period (i.e., 5-8h, Fig. 2B) and 3/ the peak of activity during the first 4 hours of the last dark period (i.e., 53-56h, Fig. 2B).

### Abnormal explorative behavior in FXS mice

When analyzing the distance traveled per hour in FXS mice, an initial hyperlocomotive phenotype is observed, which gradually normalizes over time. This heightened locomotion in FXS mice, in comparison to the control group, is especially pronounced during the acclimation phase in a new environment and the subsequent first period of darkness. However, during the final night spent in the same environment, both genotypes exhibit similar levels of locomotor activity (Fig. 3A). This observation is consistent with the notion of a hyperactive phenotype in FXS mice when exposed to an unfamiliar environment.

**Figure 3.**
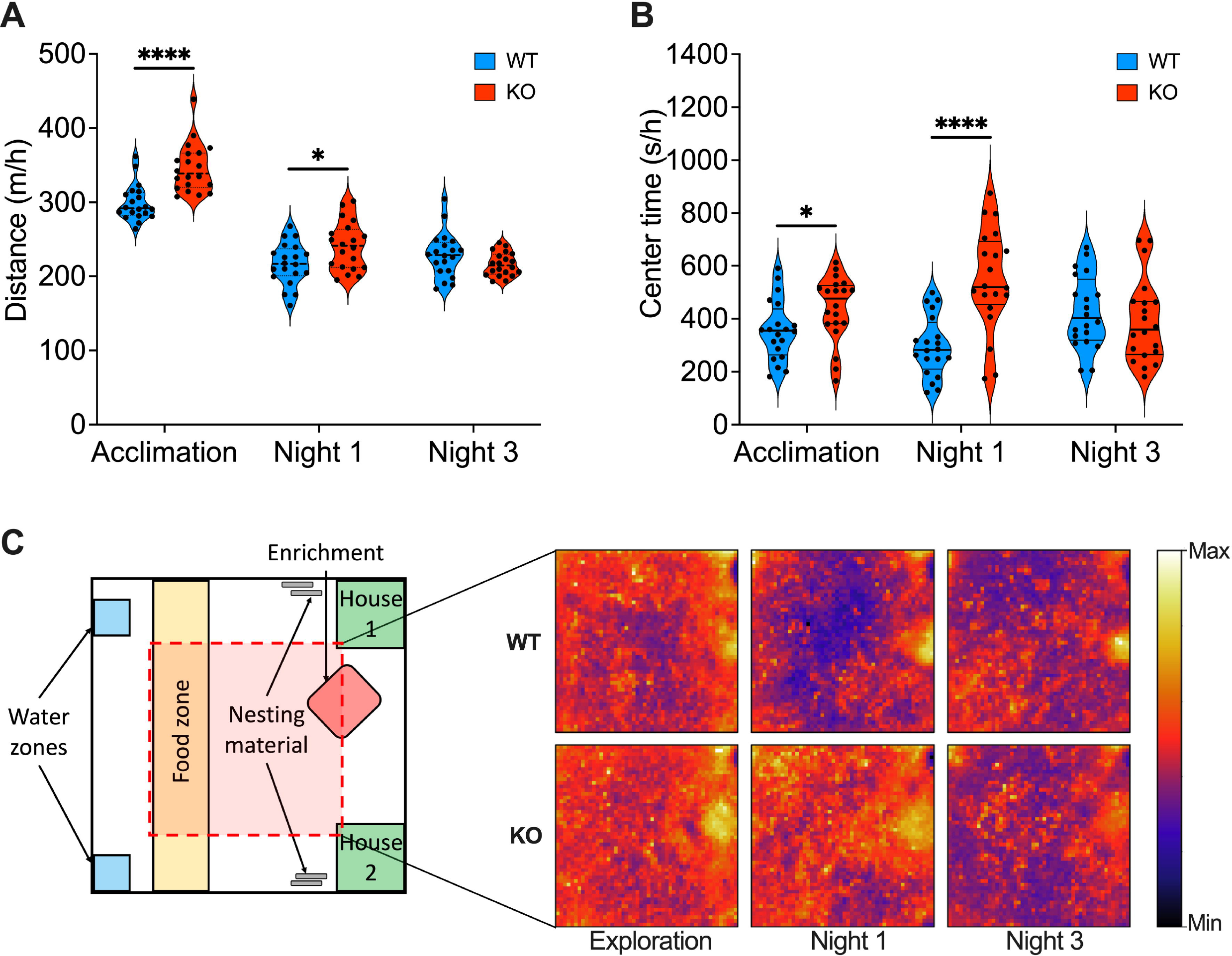
FXS mice exhibit increased locomotion and a preference for exploring the center of the arena. **A**. FXS mice display hyperactivity during the acclimation and the first dark phases. **B**. During these same phases, FXS mice spend more time in the center of the arena compared to the controls. **C**. The average time spent per hour in the center of the arena by each experimental group was transformed into a heatmap for each analyzed period. **A**-**B**. Single dot represents an individual mouse. Data are shown as violin plot with median and quartiles. Mann-Whitney U tests. ⍰= p <0.05; ⍰⍰ = p <0.01; ⍰⍰⍰ = p <0.001; ⍰⍰⍰⍰ = p <0.0001. WT group in blue, N=20; KO group in orange, N=20.

In the natural behavior of rodents, they tend to avoid open spaces to evade potential predators. The time spent in the center of the arena serves as a general indicator of the mouse’s anxiety level. In parallel with their hyperactive locomotion phenotype, FXS mice spend a higher amount of time in the center during the initial phases of the experiment, while this parameter aligns more closely with the behavior of control mice during later phases (Fig. 3B, C).

Rearing, an essential element of exploratory behavior that indicates the mouse’s search phase during its interaction with the environment, was also extracted (Fig. 4). Interestingly, evaluation of the total time spent in rearing, showed that FXS mice exhibited a deficit in the late stage (third dark period) of the experiment, but not in the early stages (acclimation or first dark period, Fig. 4C). This contrasts with the locomotor and center exploration abnormalities that were primarily evident in the early stages (Fig. 3). This deficit in rearing behavior in FXS mice during the late stage can be attributed to both a lower frequency and a shorter average duration of this behavior (Fig. 4A, B). To investigate the influence of conspecifics, rearing behavior in contact of a cage mate (Fig. 4H) and during isolated rearing (Fig. 4D) were analyzed. During the final night, both the average (Fig. 4F, J) and total (Fig. 4G, K) duration of both behaviors was reduced in FXS mice. During the first night only “rearing in contact” was reduced (Fig. 4K). Furthermore, the frequency of isolated rearing did not display statistically significant differences in any of the phases (Fig. 4E), while rearing in contact was less frequent during the third night (Fig. 4I). None of the forms of rearing exhibited alterations during the acclimation to the new environment (Fig. 4). FXS mice exhibit impaired rearing behavior, particularly in the later stages of the experiment, with more pronounced alterations observed in rearing in contact as compared to isolated rearing.

**Figure 4.**
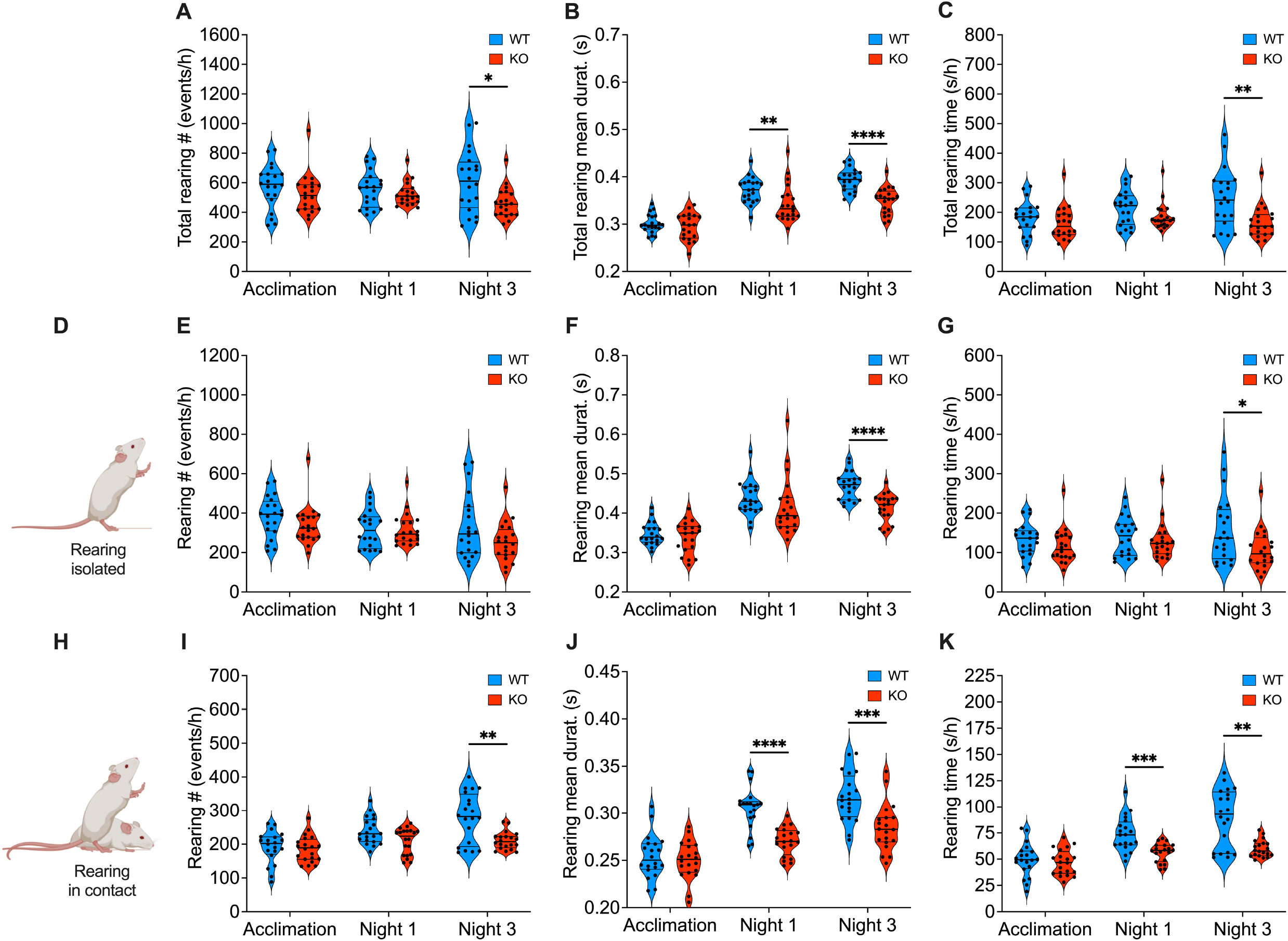
Rearing behavior in FXS mice is deficient. **A-C**. Compared to controls, FXS mice exhibit less frequent rearing during the third night (A), with shorter mean duration the first and third night (B) and less total duration the third night (C). **D**. Illustration of rearing isolated behavior. **E-G**. Rearing isolated frequency is similar between genotypes, while its average (F) and total (G) duration are shorter in KO mice than WTs during the third night. **H**. Illustration of rearing in contact behavior. **I-K**. Compared to controls, FXS mice show less frequent rearing in contact during the third night (I), with shorter average (J) and total (K) duration during the first and third night. **A-C, E-G, I-K**. Single dot represents an individual mouse. Data are shown as violin plot with median and quartiles. Mann-Whitney U tests. ⍰ = p <0.05; ⍰⍰ = p <0.01; ⍰⍰⍰ = p <0.001; ⍰⍰⍰⍰ = p <0.0001. WT group in blue, N=20; KO group in orange, N=20.

### Abnormal social interactions in FXS mice

Impairments in social interactions are a prominent characteristic of the behavioral patterns observed in ASD mouse models, including FXS^32^. The examination of continuous group behaviors over extended durations has been relatively scarce in previous research. Thus, we identified, compared, and analyzed various facets of social interaction within our mouse groups. First, we examined how FXS mice engaged in and maintained physical contact with their cage mates across various experimental phases. Surprisingly, during the acclimation to the new environment, FXS mice spent significantly more time in physical contact compared to control mice (Fig. 5A). Additionally, during the acclimation and the first dark period, we observed abnormalities in the pattern of physical contact. In these phases, FXS mice engaged in physical contact significantly more frequently but for shorter durations compared to controls (Fig. 5B, C).

**Figure 5.**
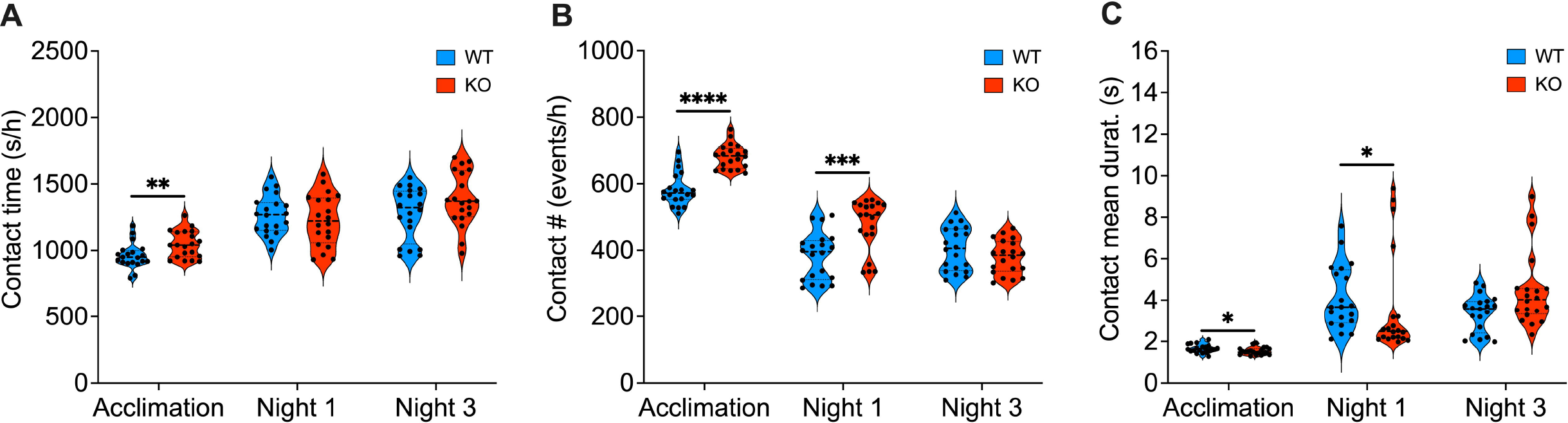
Abnormal physical contact in FXS mice. **A**. FXS mice, compared to controls, display a longer overall duration of physical contacts during the acclimation. **B, C**. Physical contacts are more frequent (B) but shorter (C) in FXS mice compared to WTs, during the acclimation and first night. **A-C**. Single dot represents an individual mouse. Data are shown as violin plot with median and quartiles. Mann-Whitney U tests. ⍰ = p <0.05; ⍰⍰ = p <0.01; ⍰⍰⍰ = p <0.001; ⍰⍰⍰⍰ = p <0.0001. WT group in blue, N=20; KO group in orange, N=20.

In summary, when placed in an unfamiliar environment, FXS mice displayed social abnormalities characterized by an increased tendency for physical contact. This contact was more frequent but less sustained compared to controls. However, as the mice became accustomed to the new environment, this behavior normalized and returned to levels similar to the control group.

Considering the observed changes in how FXS mice initiate and maintain social contact, their physical interactions were categorized according to distinct engagement mechanisms. Physical contact among mice can occur in various ways, including side-by-side contacts where two mice are positioned either facing in the same direction (S-S) (Fig. 6A) or opposite directions (S-S opposite) (Fig. 6E). During the acclimation to a new environment, FXS mice spent a cumulative amount of time engaged in S-S and S-S opposite contacts that was comparable to controls (Fig. 6B, F). However, these contacts exhibited notable qualitative differences: S-S contacts were more frequent (Fig. 6C) but of shorter duration (Fig. 6D) in FXS mice compared to WT mice. Meanwhile, S-S opposite contacts during the same period were more frequent in FXS mice (Fig. 6G) without statistically significant differences in their average duration (*p*= 0.0524) (Fig. 6H). Additionally, a shorter mean duration of S-S contacts in FXS mice was also observed during the first night (Fig. 6D).

**Figure 6.**
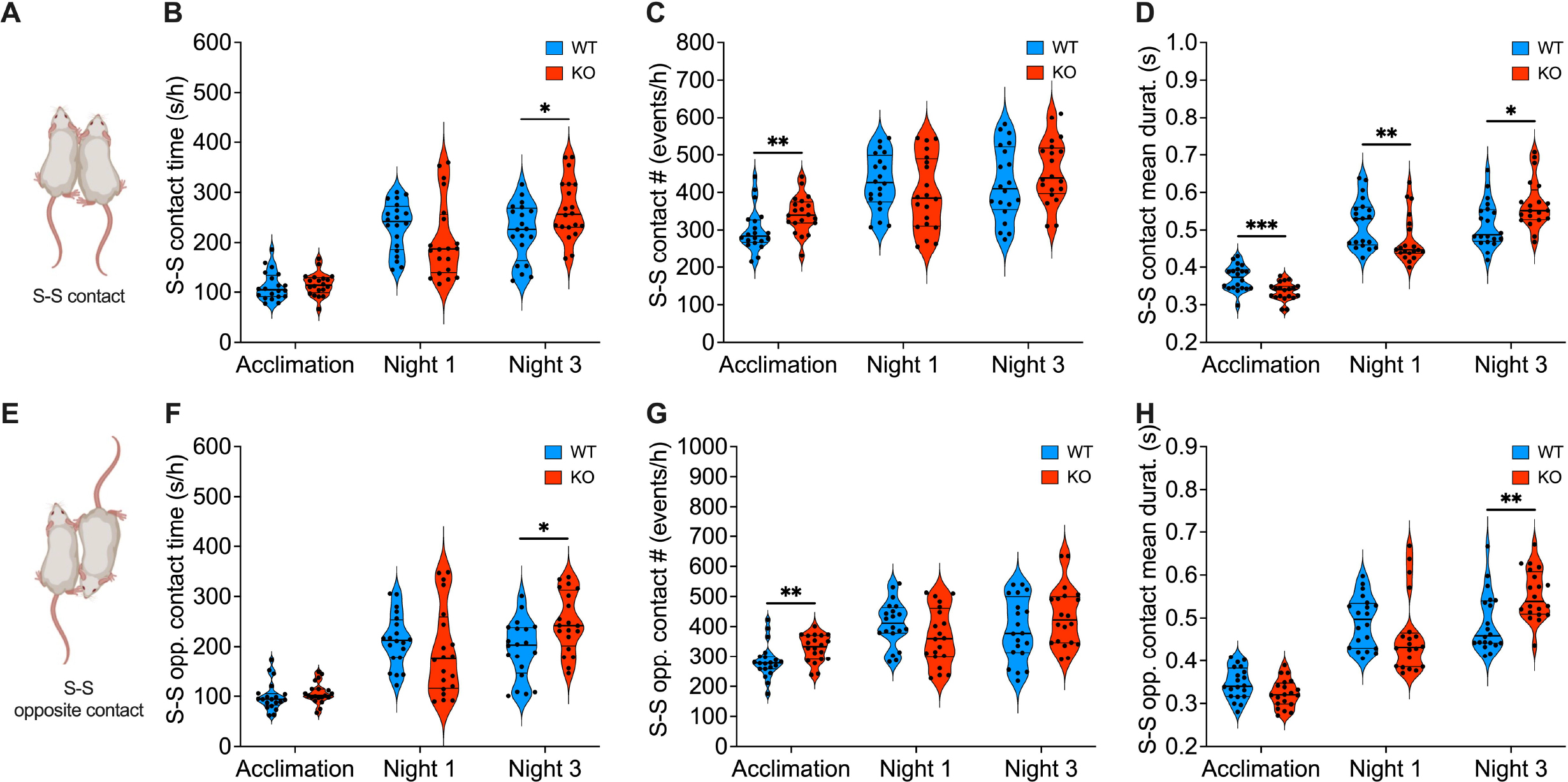
The pattern of side-by-side contacts is changed in FXS mice. **A**. Illustration of 2 mice oriented in the same direction in S-S contact. **B**. FXS mice spend more time in S-S contact than controls exclusively during the third night. **C**. FXS mice have more S-S contacts than controls exclusively during the acclimation phase. **D**. The mean duration of S-S contacts in FXS mice is shorter than that observed in controls during the acclimation and the first night, but it becomes longer during the third night. **E**. Illustration of 2 mice oriented in opposite direction in S-S opposite contact. **F**. FXS mice spend more time in S-S opposite contact than controls exclusively during the third night. **G**. FXS mice have more S-S opposite contacts than controls exclusively during the acclimation phase. **H**. The mean duration of S-S opposite contacts of FXS mice is longer than that in controls exclusively during the third night. **B-D, F-H**. Single dot represents an individual mouse. Data are shown as violin plot with median and quartiles. Mann-Whitney U tests. ⍰ = p <0.05; ⍰⍰ = p <0.01; ⍰⍰⍰ = p <0.001. WT group in blue, N=20; KO group in orange, N=20.

Conversely, during the last analyzed period (third night), FXS mice spent more time engaging in both types of contacts (Fig. 6B, F). These contacts displayed a normal frequency (Fig. 6C, G) but longer average duration compared to controls (Fig. 6D, H).

These results substantiate the notion that FXS mice exhibit nuanced anomalies in their engagement in side-by-side physical contact, with the nature of these anomalies varying based on their familiarity with the environment.

Sniffing in mice plays a crucial role as a form of communication and information gathering^33^, making it a vital component of social hierarchy and recognition^33,34^. In mice, sniffing is remarkably dynamic and is subject to variations depending on the context of their behavior. It is strongly influenced by both olfactory and non-olfactory stimuli^35^. Furthermore, mice adjust their sniffing frequency in response to different social situations, with reciprocal sniffing behavior aiding in the establishment and maintenance of social hierarchies within a group of mice^35^. In essence, sniffing behavior equips mice with the ability to navigate complex social interactions and effectively sustain their social relationships.

We examined two distinct forms of sniffing-based contacts: nose-to-anogenital (N-AG) (Fig. 7A) and nose-to-nose (N-N) (Fig. 7E). During the acclimation to the new environment, FXS mice indeed devoted more time to engaging in N-AG contact compared to control mice (Fig. 7B). These N-AG contacts were notably more frequent, while the average duration remained similar between the two groups. In contrast, during the final phase of the experiment (i.e., the third night), the total duration of N-AG interaction was higher in FXS mice than in controls (Fig. 7B) due to a longer mean duration (Fig. 7D) rather than a higher frequency of these events (Fig. 7C). FXS mice displayed similar alterations in N-N interaction as observed in N-AG interaction during the final night (Fig. 7F-H). However, during the acclimation phase, N-N interaction showed a comparable total duration between the groups (Fig. 7F), because of a higher frequency (Fig. 7G) and shorter duration (Fig. 7H) of these events in FXS mice compared to controls. Intriguingly, during the first night, FXS mice exhibited typical sniffing-based contacts (Fig. 7B-D, F-H). Usually, social interaction among mice involves a sequence of N-N and N-AG contacts (Fig. 7I). Here, FXS mice displayed abnormal frequencies of transitions from N-N to N-AG (Fig. 7J), while the reverse transitions occurred at a similar rate to controls (Fig. 7K). Specifically, compared to controls, FXS mice had fewer transitions from N-N to N-AG during the initial dark phase spent in the arena. Conversely, 48 hours later, the opposite pattern emerged, with these mice displaying a higher number of N-N to N-AG transitions (Fig. 7J).

**Figure 7.**
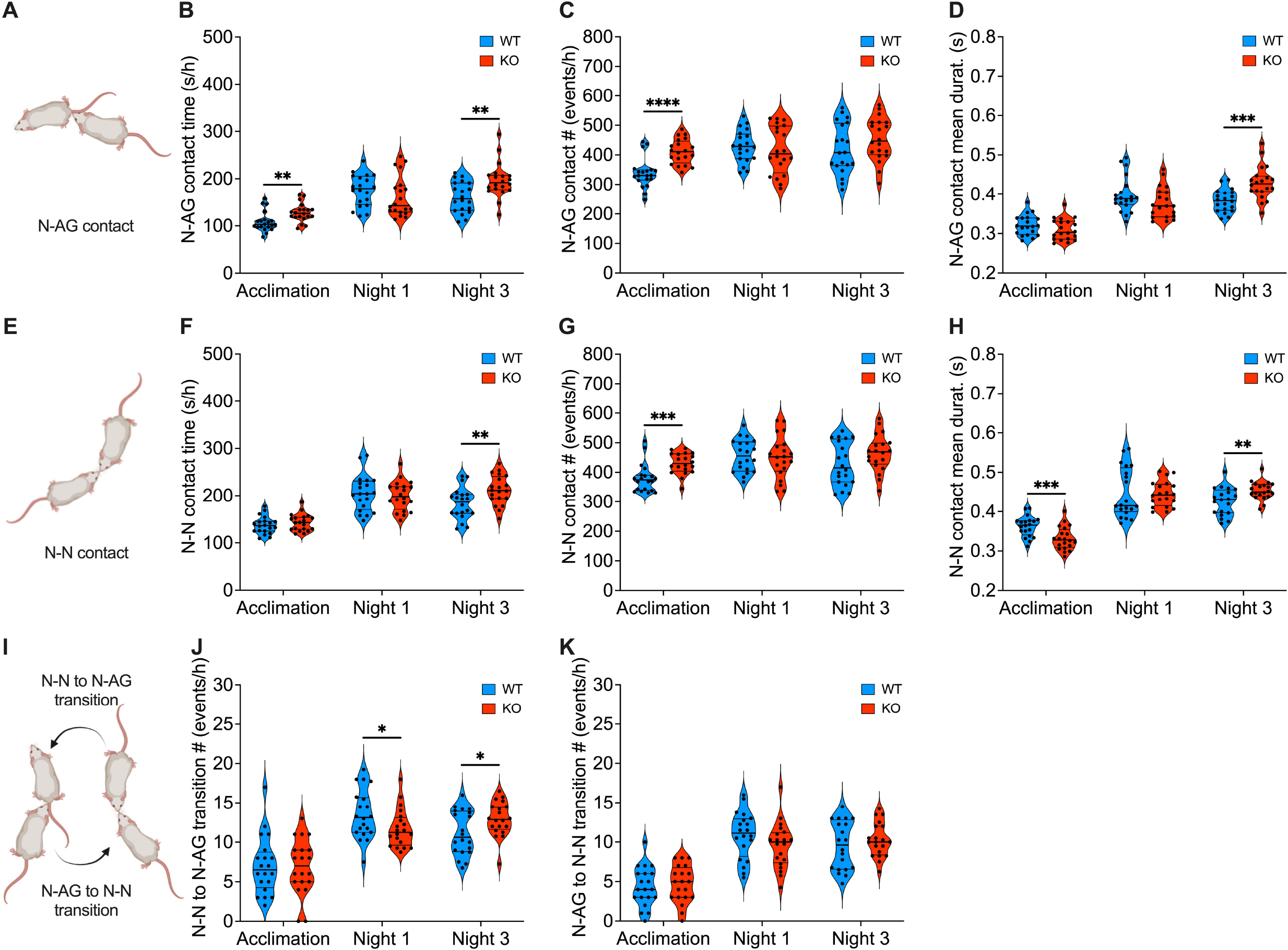
Social sniffing sequences are altered in FXS mice. **A**. Illustration of N-AG contact in mice. **B**. FXS mice spend more time in N-AG contact during the acclimation and the third night. **C**. N-AG contact frequency is higher for FXS mice during the acclimation. **D**. The mean duration of N-AG contact of FXS is longer compared to controls during the third night. **E**. Illustration of N-N contact in mice. **F**. FXS mice spend more time in N-N contact during the third night. **G**. N-N contact frequency is higher for FXS mice during the acclimation. **H**. The mean duration of N-N contact of FXS mice is shorter than controls during the acclimation, but it becomes longer during the third night. **I**. Illustration of N-N to N-AG contact transition and vice versa in mice. **J**. FXS mice display more and less N-N to N-AG transitions than controls during the first and third night respectively. **K**. N-AG to N-N transitions were comparable between groups. **B-D, F-H, J, K**. Single dot represents an individual mouse. Data are shown as violin plot with median and quartiles. Mann-Whitney U tests. ⍰ = p <0.05; ⍰⍰ = p <0.01; ⍰⍰⍰ = p <0.001; ⍰⍰⍰⍰ = p <0.0001. WT group in blue, N=20; KO group in orange, N=20.

In summary, the social interactions of *Fmr1* KO mice exhibit phase-dependent alterations in the initiation and maintenance of sniffing-based contacts. When two moving mice engage in an N-AG interaction, it is referred to as a “train” contact (Fig. 8A). This behavior combines elements of social interaction (N-AG sniffing) and exploration (movement). Throughout the analyzed periods, this train movement primarily involves only two mice at a time (Fig. 8B)

**Figure 8.**
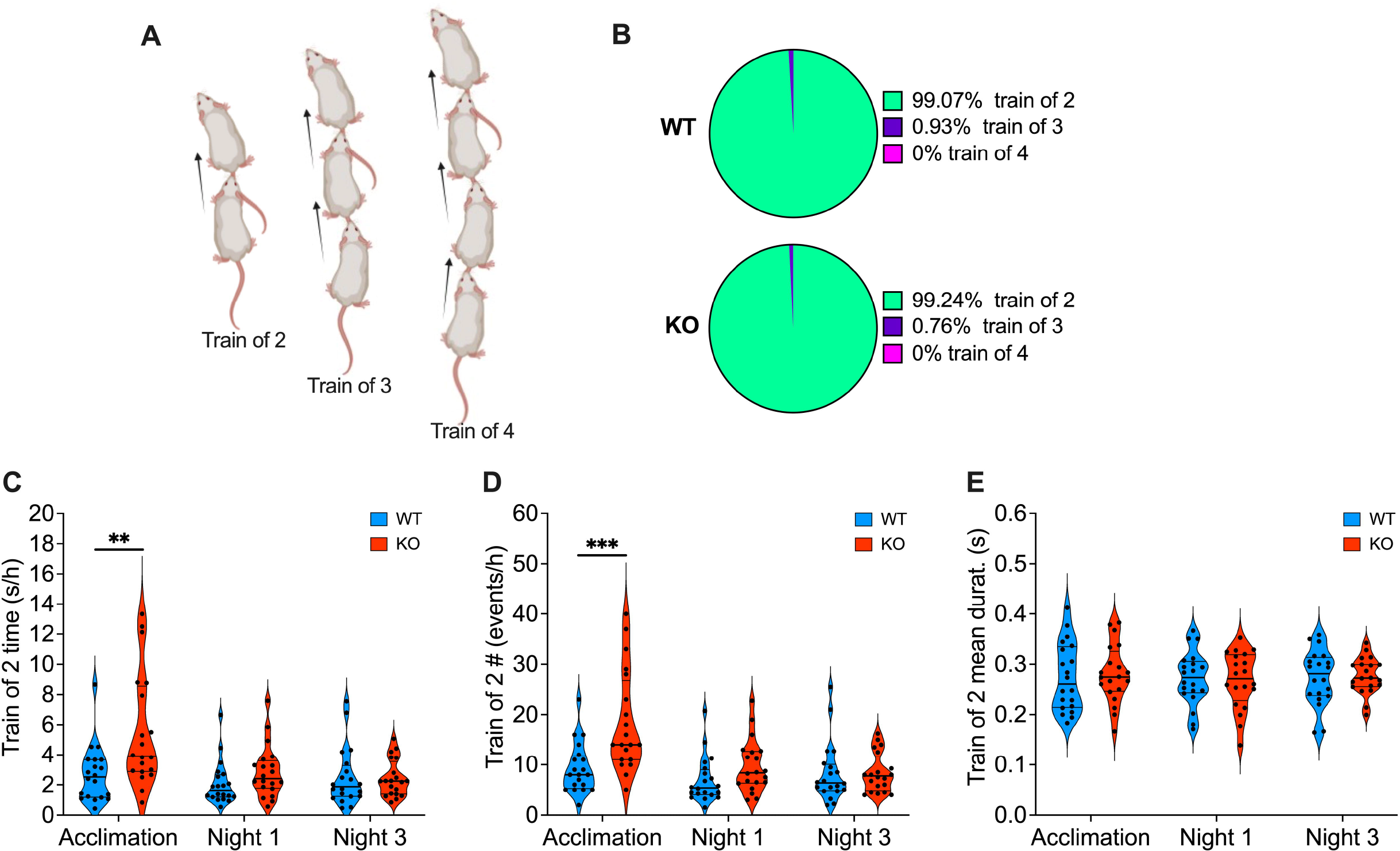
FXS mice move more in “train” when exploring a new environment. **A**. Train contacts illustration with 2, 3 or 4 mice involved. **B**. In both genotypes, train contacts involve almost always 2 mice at a time. Pie graphs: percentages of train of 2, 3 or 4 mice, in Fmr1 WT and KO groups (total time spent in train behavior over the three analyzed periods by each group). **C-E**. FXS mice, compared to controls, spend more time moving in a train of 2 during the acclimation phase (C), where they engage more frequently in this type of contact (D) but with the same mean duration (E). Single dot represents an individual mouse. Data are shown as violin plot with median and quartiles. Mann-Whitney U tests. ⍰⍰ = p <0.01; ⍰⍰⍰ = p <0.001. WT group in blue, N=20; KO group in orange, N=20.

In contrast to the overall N-AG contact observations, the “train of 2” behavior shows a longer total duration in FXS mice compared to controls, specifically during the acclimation to the new environment (Fig. 8C). This anomaly can be attributed to a higher frequency of engagement in this contact (Fig. 8D), while the average duration remains the same between the two groups (Fig. 8E).

These results suggest that in an unfamiliar environment, FXS mice tend to move more closely behind one another compared to controls.

### The familiarity of the environment determines whether FXS mice move in groups or independently

The anomalies observed so far support the hypothesis that FXS mice tend to favor exploring a new environment in groups. To test this hypothesis, we assessed their inclination toward either moving in contact with another mouse or moving alone during different phases of the experiment (Fig. 9). During the acclimation phase and the first dark period in the new environment, FXS mice indeed dedicated more time to moving while in contact with a cage mate compared to controls. However, during the third dark period, this behavior returned to normal (Fig. 9B), and no significant differences were noted in the time spent not moving in contact with another mouse (Fig. 9D). Conversely, the cumulative time spent moving alone was higher in FXS mice only during the first night (Fig. 9F), while the time spent not moving alone was significantly lower in FXS mice compared to controls, particularly during the acclimation and the last night (Fig. 9H). FXS mice displayed a stronger preference than the control group for moving in contact with others, especially during the acclimation to the new environment (Fig. 9I). In contrast, when it comes to phases of non-movement, KO mice exhibited a greater preference for physical contact only during the last dark period (Fig. 9J).

**Figure 9.**
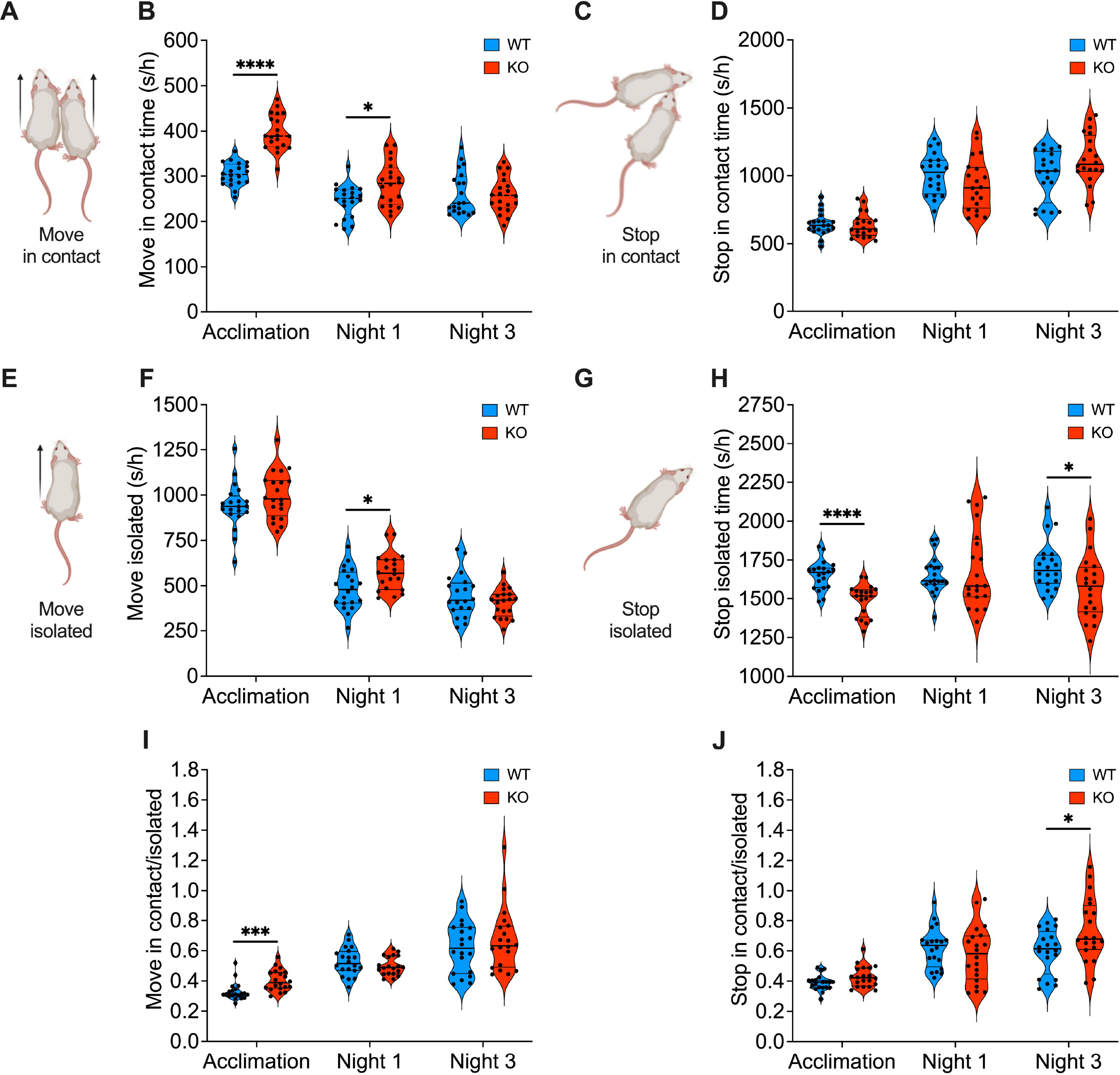
FXS mice show elevated preference for social contact while moving or not, depending on the familiarity with the environment. **A**. Illustration of two mice moving in contact. **B**. FXS mice spend more time moving in contact than controls specifically during acclimation and first night. **C**. Illustration of two stationary mice in contact. **D**. The time spent in contact during non-moving phases is similar between groups. **E**. Illustration of a mouse moving alone. **F**. FXS mice spend more time than WTs in moving isolated during the first dark period. **G**. Illustration of a stationary isolated mouse. **H**. FXS mice spend less time alone when they do not move during the acclimation and last dark period. **I, J**. FXS mice, compared to controls, have higher preference for contact while moving during acclimation (I) and, conversely, while stationary during the last night (J). **B, D, F, H, I, J**. Single dot represents an individual mouse. Data are shown as violin plot with median and quartiles. Mann-Whitney U tests. ⍰ = p <0.05; ⍰⍰⍰ = p <0.001; ⍰⍰⍰⍰ = p <0.0001. WT group in blue, N=20; KO group in orange, N=20.

These findings indicate that in an unfamiliar environment, FXS mice are less likely to explore alone compared to controls. As the environment becomes more familiar, they are more willing to be alone while exploring, but this preference disappears when they stop.

### Group dynamics are different in FXS mice

Group dynamics are influenced by the individual social skills of their members, and changes in the social interaction patterns of FXS mice can have an impact on these dynamics. Consequently, an analysis of group formation parameters was carried out. In a group consisting of four individuals of the same sex, mice from both genotypes tended to spend the majority of their social contact time in pairs (approximately 69%), followed by groups of three (around 29%), with instances of being in groups of four being quite rare (below 2%) (Fig. 10A, B). Therefore, the analysis primarily focused on events involving the formation of groups composed of two or three mice. In comparison to the control group, FXS mice spent an equivalent amount of time in pairs throughout all the phases analyzed (Fig. 10C). However, during the acclimation phase, they allocated more time to being in groups of three (Fig. 10F). Furthermore, the formation of groups of both sizes occurred more frequently (Fig. 10D, G) and had shorter durations (Fig. 10E, H) among FXS mice compared to controls, but only during the acclimation to the new environment and the first dark period spent within it.

**Figure 10.**
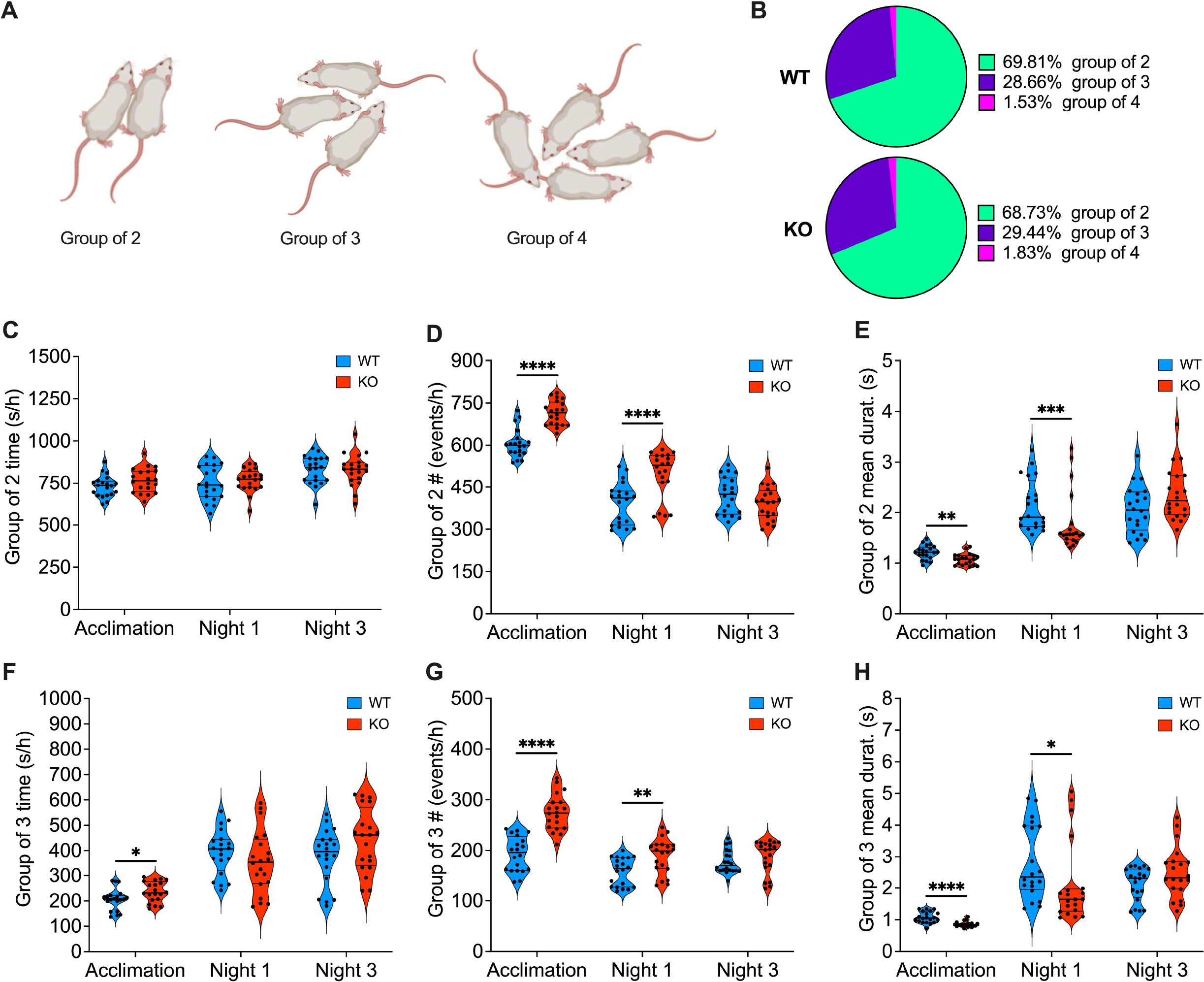
FXS mice show abnormal group dynamics depending on the familiarity of the environment. **A**. Illustration of groups of 2, 3 or 4 mice. **B**. Mice of both genotypes predominantly engage in social interactions in pairs (∼69%), followed by groups of three (∼29%), and rarely in groups of four (below 2%). Pie graphs: percentages of group of 2, 3 or 4 mice for Fmr1 WT and KO mice (total time spent in group over the three analyzed periods by each genotype). **C-E**. The total time spent in pairs does not vary between genotypes (C), although FXS mice form pairs more frequently (D) and with shorter durations (E) during acclimation and the first night. **F-H**. During acclimation, groups of 3 FXS mice lasted longer overall, were more numerous (G), but each individual mouse had shorter durations (H). **C-H**. Single dot represents an individual mouse. Data are shown as violin plot with median and quartiles. Mann-Whitney U tests. ⍰ = p <0.05; ⍰⍰ = p <0.05; ⍰⍰⍰ = p <0.001; ⍰⍰⍰⍰ = p <0.0001. WT group in blue, N=20; KO group in orange, N=20.

In summary, in a novel environment, FXS mice display a significantly higher frequency of forming and dissolving social groups, which happens more rapidly than in controls. This phenotype normalizes when the environment becomes familiar.

In conclusion, in line with most of the previously presented data, the PCA conducted on the exploratory and social phenotype in FXS mice (see Fig. 11) corroborates that the most notable distinctions between genotypes were evident when the mice were initially exposed to an unfamiliar environment (Fig. 11A). Intriguingly, the phenotype progressively returned to levels like those of the control group as the mice became more accustomed to their surroundings over time (Fig. 11C).

**Figure 11.**
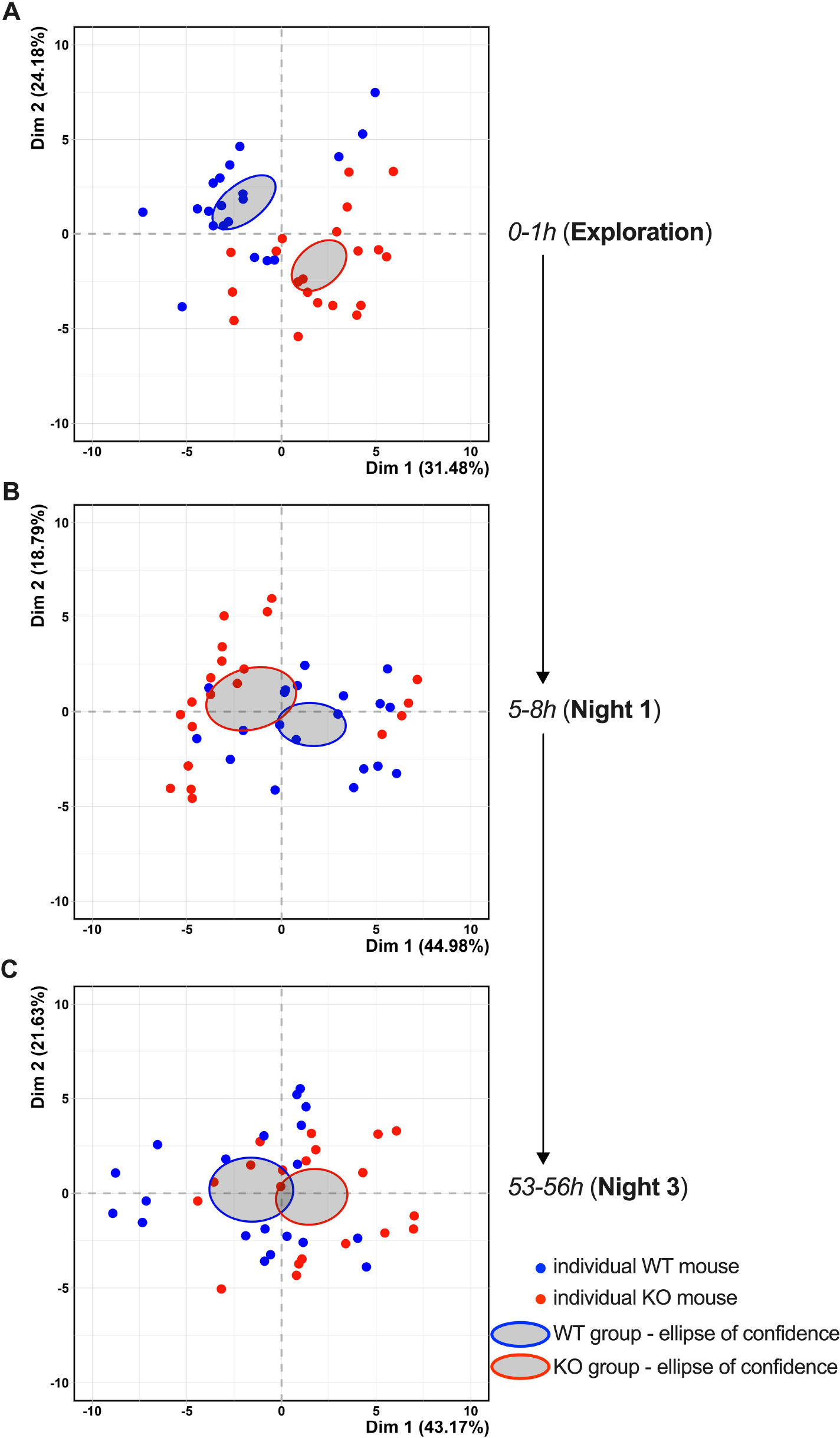
FXS mice display atypical behavior upon initial exposure to an unfamiliar environment that gradually normalizes with time. **A-C**. Principal component analysis (PCA) of exploratory and social behavior shows an atypical phenotype of FXS mice during the first hour in an unfamiliar environment (A) that gradually normalizes in the later analyzed phases (B and C). PCA was performed on the behavior recorded during 0-1h, 5-8h, 53-56h time intervals. Behaviors included in PCA are: distance traveled, center time, rearing, total contacts, S-S contacts, N-AG contact, N-N contacts, train of 2, move in contact, move isolated, stop in contact, stop isolated, group of 2 and group of 3. Single dot represents an individual mouse. Ellipses represent the barycenter and confidence interval per group. Data are shown as PCA of individuals, dimension 1 and 2 percentages of variance are displayed in graphs. WT group in blue, N=20; KO group in orange, N=20

## Discussion

In this study, we compared the continuous, unsupervised, and spontaneous social interactions between groups of WT and FXS mice to understand the dynamics of social behavior in FXS. The findings reveal complex abnormalities in the exploratory and social behavior of FXS mice, with the most significant differences occurring during their initial adjustment to a new environment. Over time, these altered behaviors gradually decreased and returned to normal. This dynamic FXS phenotype is characterized by heightened activity, a greater inclination to explore exposed areas, increased interest in social interaction, and a distinctive pattern of social interactions.

The findings indicate that groups of FXS and WT mice exhibited a comparable day/night rhythm in locomotor activity, consistent with previous research on individually housed FXS mice (e.g., a 25-hour experiment by Pietropaolo et al., 2011^36^, and a 16-day observation by Bonasera et al., 2017^37^). While sleep problems have been observed in humans with FXS^38,39^, and alterations in circadian clock-related parameters have been noted in the Drosophila model of FXS^40,41^, it appears that these issues do not have apparent effects on the locomotor activity rhythms of mice in day/night cycles. In this study, we observed that FXS mice displayed hyperactivity during the initial phases of the experiment when they were active, but not during the later stages. This hyperactive behavior appeared to be induced by exposure to an unfamiliar environment. It is worth noting that individuals with FXS often exhibit symptoms of hyperactivity^42^.

Consistent evidence points to a hyperactive locomotor phenotype in FXS mice when assessed using conventional tests like the open field^43–52^. However, these tests, which usually last for 30 to 120 minutes, are not optimal for gaining a comprehensive understanding of the novelty-dependent aspects of this behavior. Consistent with our findings, a study documented a hypoactive locomotor phenotype in FXS mice for 16 days after a 5-day habituation period in their familiar environment^37^. Thus, FXS mice may transition between hyperactive and hypoactive phenotypes depending on the level of environmental familiarity.

Individuals with FXS are frequently diagnosed with anxiety disorders^12^. Convergent evidence from mouse models of FXS demonstrates anxiety-related alterations in test settings such as the open field^44,48,49,52,53^ and the elevated plus/zero maze^20,43,44,53–55^. We observed altered anxiety phenotypes in FXS groups depending on the level of environmental familiarity. FXS mice spent more time in the center of the arena during the early stages of the experiment but not in the late stages, consistent with the literature

Social deficits are a common feature of both ASD and FXS^56^. In our FXS groups, FXS mice exhibited an increased total time spent in physical contact with a cage mate during their initial adjustment to the new environment, but this behavior normalized to levels like controls in later stages. Specifically, during the acclimation phase and the initial active period (i.e., the first dark period), social contacts were more frequent but of shorter average duration compared to controls. We hypothesize that this behavior could reflect an elevated interest or need for social contact in response to discomfort. Noteworthy, the increase in social contacts coincided with the presence of hyperactive and altered anxiety phenotypes, implying a potential influence of these traits on social skills. This observation aligns with existing research that highlights the significant impact of hyperactivity^57^ and anxiety^58^ on social functioning in individuals with FXS. Furthermore, the pattern of ‘frequent approaching followed by rapid disengagement’ resembles an approach-avoidance conflict, as indicated by several clinical studies. Despite heightened social anxiety and avoidance tendencies, individuals with FXS are also reported to exhibit behaviors suggesting their willingness or desire to interact with others^8,58–62^.

In this comprehensive examination of the FXS phenotype in groups of mice, the most notable disparities between genotypes emerged during the initial exposure to an unfamiliar environment. Subsequently, selected phenotypes exhibited a gradual normalization, converging toward levels comparable to those of the control group as the mice became more acclimated to their surroundings. This observation aligns with heightened sensitivity traits, including increased arousal^14^, sensory hypersensitivity^63^, and reduced predictive capabilities^64^, often observed in individuals with FXS. These traits lead to an enhanced perception of novelty in environmental stimuli. Consequently, encountering uncomfortable situations, such as unfamiliar environments, can potentially trigger or exacerbate behavioral abnormalities in FXS individuals. Understanding the neural mechanisms underlying this dynamic phenomenon may hold significant implications for the development of innovative therapeutic strategies, both in terms of behavioral interventions and pharmacological approaches.

## Abbreviations

FXS: Fragile X Syndrome
ID: intellectual disability
ASD: autism spectrum disorders
FMRP: Fragile X Messenger Ribonucleoprotein 1
WT: wild type
KO: knockout
LMT: Live Mouse Tracker
PCA: Principal Component Analysis
S-S: side-by-side
N-AG: nose-to-anogenital
N-N: nose-to-nose

## Declarations

### Ethics statement

Animals were treated in compliance with the European Communities Council Directive (86/609/EEC) and the United States National Institutes of Health Guide for the care and use of laboratory animals. The French Ethical committee authorized this project (APAFIS#34573-202201071122121v3).

### Availability of data and materials

All data reported in this paper will be shared by the lead contact upon request. Any additional information required to reanalyze the data reported in this paper is available from the lead contact upon request.

### Authors’ contributions

GG: Conceptualization, Data curation, Formal analysis, Validation, Writing-review and editing.

BS: Conceptualization, Data curation, Formal analysis, Validation. OL: Validation, Methodology.

PC: Conceptualization, Supervision, Methodology, Writing-review and editing. OJM: Conceptualization, Supervision, Funding acquisition, Methodology, Project administration, Writing-original draft, review and editing.

## Acknowledgements

The authors are grateful to the Chavis-Manzoni team members for helpful discussions.

